# Neural correlates in basolateral and central amygdala during reward seeking in the face of punishment

**DOI:** 10.1101/2025.03.26.645435

**Authors:** Angelica Minier-Toribio, Brian Sadacca, Emmanuelle Corruble, Denis J. David, Yavin Shaham, Geoffrey Schoenbaum, Yann Pelloux

## Abstract

The inability to suppress actions despite adverse consequences is a hallmark of compulsive behaviors and is amygdala dependent. To study the amygdala’s role in responding despite adverse consequences, we compared single-unit activity in the basolateral (BLA) or central (CeA) amygdala before and after reward-seeking under punishment threat.

Rats started each trial by pressing an initial lever, which triggered an outcome-specific 5-s auditory cue (white noise, pure tone, or clicker), signaling one of three reinforcement conditions: 100% sucrose reward, 20% reward/80% omission, or 80% reward/20% footshock punishment. Next, the active and inactive levers were extended, and pressing on the active lever terminated the auditory cue and triggered an outcome-specific 1-s visual cue predicting either reward, reward omission, or shock. After training, we implanted eight tetrodes in the BLA (n=7) or CeA (n=5) and recorded single-unit activity of ∼100 neurons per region during task performance.

CeA neurons, and to a lesser extent BLA neurons, responded differently to the distinct auditory cues and outcomes. The discrimination between conditions partially explained the capacity of neuronal activity to predict the latency to lever press to complete the trial. The failure to suppress reward-seeking behavior in the face of punishment coincided with the reactivation of reward-seeking-sensitive and the loss of inhibition of punishment-sensitive neuronal populations.

In conclusion, we found that after extended training, opposing populations of activated and inhibited neurons in CeA, and to a lesser extent in BLA, control completion of reward-seeking despite punishment.

## Introduction

Persistent responding despite adverse consequences is a key feature of compulsive disorders, including drug addiction. Different preclinical rodent models have operationalized compulsive drug use by coupling drug-related behaviors with punishment. For instance, extended exposure to cocaine or alcohol results in a subpopulation of rats that maintain drug self-administration despite intermittent footshock punishment (1-7). Lesion of the basolateral amygdala (BLA) and pharmacological inactivation of the central nucleus of the amygdala (CeA) partially mimic this phenotype in rats with a limited history of cocaine self-administration (8,9). Additionally, BLA inactivation prevents suppression of food seeking in the face of punishment (10-15). Pharmacological inactivation of CeA or BLA also reinstates drug seeking after punishment-induced suppression of drug self-administration (16,6) These studies suggest an active role of BLA and CeA neurons in suppressing punished responses. However, they contradict recent findings of increased BLA activity during lever pressing under punishment when suppression fails (17). Furthermore, CeA inactivation using the GABAb agonist baclofen has been shown to reduce punishment-resistant alcohol seeking in rats with extended alcohol reinforcement histories (18). Together, no clear picture has emerged on the role of BLA and CeA neuronal activity in the control of reward seeking under punishment.

We investigated the role of the amygdala in punishment-induced suppression of operant food self-administration by recording single-unit activity within the BLA and CeA. We used a behavioral procedure designed to dissociate deliberative and executive processes occurring before the outcome from the subsequent reevaluation of the action. Our main goal was to determine whether neural responses within the amygdala could discriminate between cues and outcomes based on their valence and predict the subsequent behavioral response.

## Material and Methods

### Subjects

We initially maintained 14 male Sprague–Dawley rats (Charles River), weighing 250–350 g, by pairs under a reverse 12:12 h light/dark cycle (lights off at 8:00 a.m.) with food and water freely available. One week after arrival we restricted the rats to 80% of their daily food intake to ensure stable performance. Experiments complied the National Institutes of Health Guide for the Care and Use of Laboratory Animals (8th edition) under the protocol (#15-BNRB-189) approved by the local Animal Care and Use Committee.

### Apparatus

We shielded sound-attenuating cabinets of Med Associates operant chambers (29.5 × 32.5 × 23.5 cm; Med Associates, Georgia, VT) with aluminum foil connected to a copper solid-state ground. The chamber floor consisted of 1 cm spaced metal bars, each connected to a computer-controlled switch (DB9 A/B Network Switch, electrostandards laboratories), that alternated between a shock generator (Med Associates) and the ground. One side of the chamber featured the active and inactive retractable levers, 8-8.5 cm above the grid floor, 12 cm apart; each surmounted by a cue light (2.5 W, 24 V). In between, a recessed food magazine (3.8 cm^2^ and 5.5 cm from the grid floor) fitted with infrared beams to detect magazine entries (nose-pokes) connected an external pellet dispenser. On the opposite wall, a third lever (initiation lever) stood centrally 8-8.5 cm above the grid floor, surmounted by a loudspeaker and a houselight (2.5 W, 24 V).

### Experimental design and procedure

#### Phase 1: Magazine training

Sessions started with the onset of the houselight. Rats collected one 45-mg food pellet (AIN-76A, TestDiet) from the magazine port, delivered every 60 s during a 1-hour session over 1-2 days.

#### Phase 2: Instrumental training

Sessions began with the houselight onset and the presentation of the active and inactive levers (left or right —counterbalanced across rats). Pressing on the active lever retracted the levers and triggered the delivery of a food pellet in the magazine port. We recorded inactive lever presses but they had no consequences. A 10-s timeout after reward delivery preceded the subsequent trials. Training consisted of 5 daily 1-h sessions until stable performance.

#### Phase 3: Training on 100% rewarded trials

Sessions began with the houselight onset and insertion of the initiation lever. Pressing the lever triggered its retraction and the onset of a 65-dB auditory cue, being either a white noise, a clicking sound (10 Hz) or a pure tone (8 kHz) —counterbalanced across rats. We chose the frequency and intensity of the auditory cues based on Kelly and Masterton (19). Five seconds later, the active and inactive levers extended simultaneously. Pressing on the active lever retracted both levers, silenced the auditory cue, and initiated a continuous 1-s light cue above the active lever. The light cue offset then triggered reward delivery into the magazine. After a 10-s time-out period, the presentation of the initiation lever signaled the start of a new trial. Sessions ended once rats completed 100 trials, or 60 min elapsed. At the end of the fifth session, all rats reliably self-initiated trials.

#### Phase 4: Training under intertwined contrasting trials procedure

After houselight onset (Figure 1A) and initiation lever insertion, pressing the lever triggered its retraction and the onset of 65dB auditory cues: either a pure tone (8 KHz), a clicker (10 Hz), or a white noise. Each auditory cue signaled a different contingency: 1) “reward”, 2) “reward/omission” or 3) “reward/shock” (counterbalanced across rats). The auditory cue onset triggered the presentation of the active and inactive levers 5-s later. Pressing on the active lever turned off the auditory cue, retracted the levers, and turned on a visual light cue for 1-s. This cue was either: : 1) a continuous light above the active lever signaling impending food pellet delivery upon its offset, 2) a continuous light above the inactive lever signaling the omission of reward upon its termination, or 3) a flashing light above the active lever signaling impending footshock delivery upon offset. After a 10-s time-out period, initiation lever presentation marked the start of a new trial.

**Figure 1.**
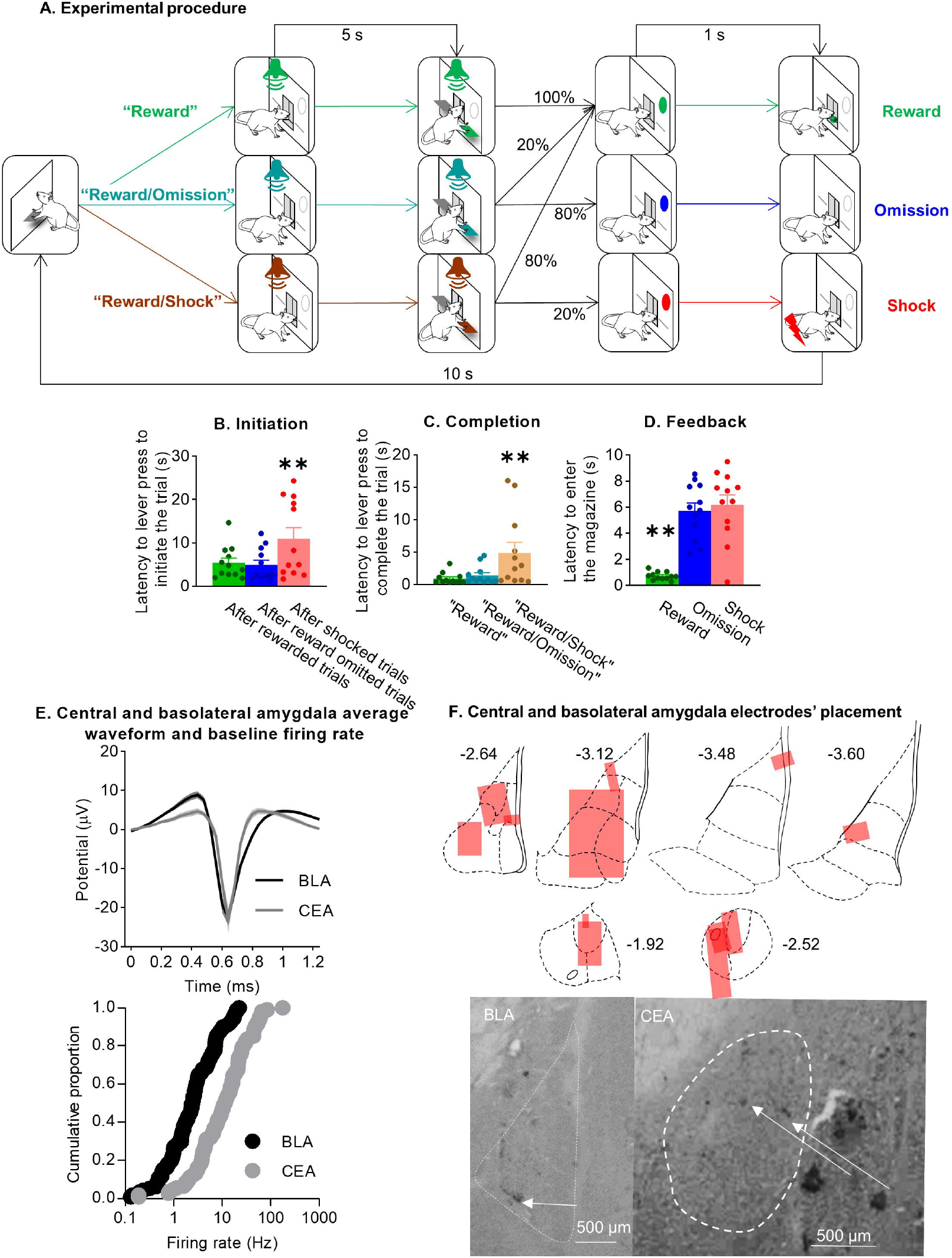
<u>(**A**) Experimental procedure</u>. (**B**) Mean ± SEM latency (in seconds) to press on the initiation lever. This sets off an auditory cue (white noise, pure tone, or clicker) that signaled the experimental condition (“reward”, “reward/omission”, “reward/shock”), followed 5 s later, by insertion of the active and inactive levers (n=12). (**C**) Mean ± SEM latency (in seconds) to press the active lever to complete the trial. This sets off a visual cue (feedback) that signals the impending outcome 1 s later (reward, omission or shock) (n=12) (**D**) Mean ± SEM latency to enter the magazine after outcome presentation (n=12). For panel (B)-(D): * Significantly different from the other two conditions Benjamini-Hochberg corrected Fisher’s LSD, p<0.05. (**E**) <u>Top panel</u>: Mean ± SEM waveform of the action potential of the BLA (black dots) and CeA (grey dots). <u>Bottom panel</u>: Cumulative frequency distribution of the mean firing rate 1 s before each trial for BLA (black dots) and CeA (grey dots) neurons. (**F**) CeA and BLA electrodes’ placement. Panels 1 and 3: schematic representations of the area of recording for the CeA (n=5) and BLA (n=7) on standardized sections of the rat brain. The boxes represent the area from which neurons were recorded in each rat using the moveable tetrode arrays. The numbers adjoining each section refer to distances from bregma [adapted from (Paxinos and Watson, 2008)]. <u>Panels 2 and 4</u>: pictures from a representative implantation into the CeA and BLA, respectively. The dashed lines delineate the targeted brain regions and arrows point the tip of the electrodes.

Each session randomly intertwined 20 “reward” trials, 40 “reward/omission” and 40 “reward/shock” trials on average. In the “reward” trials, active lever pressings always resulted in reward (100% reward) while only 20% of the pressings led to reward delivery in the “reward/omission” trials (20% reward or 80% omission). In the “reward/shock” trials, 80% active lever pressing resulted in reward delivery, and 20% led to a mild footshock (80% reward or 20% footshock).

Sessions ended once rats completed 100 trials or 1 hour elapsed. We measured 1) the latency to initiate the trial as the duration from lever presentation to lever press on the initiation lever, 2) the latency to complete the trial as the duration from the presentation of the active and inactive levers to active lever press, and 3) the latency to enter the magazine (nosepoke) from the onset of the visual cue.

To optimize auditory cue discrimination, we individually adjusted footshock intensity. We progressively increased from 0.2 mA by 0.05 mA every five days until rats performed fewer than 100 trials during the 1-h session. We then progressively reduced the intensity by 0.05 mA until rats 1) showed a 3-fold difference in reward vs. reward/shock trials of lever press latencies, and 2) completed the 100 trials in less than an hour. Rats completed training in 16 sessions for phases 1-3 and 45±3 sessions for phase 4, after which we implanted recording electrodes.

### Surgery for electrode implantation

Electrodes consisted of eight tetrodes drivable bundles. Each tetrode consisted of four twisted polyimide coated Nickel chrome wires of 25-μm-diameter (Stablohm 675; California Fine Wire). The day before implantation, we trimmed each electrode tip with surgical scissors and gold-plated to achieve an impedance of approximately 300 kΩ (20). We stereotaxically implanted the electrodes in isoflurane-anesthetized rats (5% induction; 2–3% maintenance, coordinates: BLA: anterior-posterior (AP): -2.5 mm and mediolateral (ML): 5.3 mm from bregma and dorsoventral (DV) -6.8 mm from dura (n=12). CeA: AP: -2.3 mm, ML: 4.0, DV -6.8 mm, based on (21) and our previous study (16). We anchored the electrodes to the skull with jeweler’s screws and dental cement.

After one week of recovery, rats returned to the task for one week without recording. Two rats died before we performed any recording. We recorded 1-h daily sessions at a rate of 25 kilo samples per second. We connected the electrode to a headstage (Intantech RHD2132 amplifier board), connected to an OpenEphys interface via a commutator to ensure free movement of the rats in the operant chambers. The interface also connected a TTL device whose signal was controlled by the computer operating the Med Associates interface, ensuring synchronization of the recording with events (lever presses and nosepokes) generated by the Med Associates interface. The OpenEphys interface connected a second computer that recorded the raw signals. We used the Klustaview suite for post-processing of spike detection and spike sorting of the raw signal. We then performed all analyses using Matlab®. We moved the drivable electrode ∼40 μm every third day until the electrode was located dorsal-ventral approximately -8 mm from the dura.

At the end of the study, we subjected rats to an overdose of isoflurane, marked the final electrode position by passing current through electrodes, removed their brains and stored them in 10% formalin. We sliced the brains into 75 μm sections using a Leica Microsystems cryostat and stained the sections with cresyl violet to verify the placement of the electrodes under a light microscope.

### Statistical analysis

#### Behavior

Separate one-way repeated measures analysis of variance (ANOVA) with experimental condition (reward, omission, shock) as the within-subjects factor.compared: 1) the latency to initiate trials following a food pellet reward, reward omission (omission), or shock in the preceding trial; 2) the latency to complete trials under the conditions of “reward,” “reward/omission,” and “reward/shock”; and 3) the latency to nosepoke while visual cues indicated the impending reward, omission, or shock. We followed up on significant effects using a Benjamini-Hochberg corrected Fisher’s PLSD for false discovery rate multiple-comparison posthoc test.

#### Neural activity

To account for heterogeneous neuronal firing rates, we normalized the 100 ms binned firing rate to a z-score using baseline activity. The baseline activity’s average and variance were calculated from the firing rate over the second preceding the start of the different trials. We also used non-parametric statistical tests for most analyses, with parametric methods employed only when necessary to evaluate interactions between factors.

Two-way repeated measures ANOVAs (time x conditions) assessed CeA and BLA neuronal firing rate changes following auditory/visual stimuli. We examine activity during trial completion and initiation by averaging z-scored activity across defined epochs, and compared using Friedman tests (overall population) or ANOVAs (subpopulations), followed by Benjamini-Hochberg post-hoc tests.

To stratify neurons, we compared individual activity across trial types using Kruskal-Wallis tests to assess their condition-discrimination, and correlated it with the log-transformed lever press latencies using Spearman’s correlations to assess their ability to predict behavior.

## Results

### Behavior

One-way repeated measures ANOVA demonstrated that rats successfully discriminated between the auditory cues [F(2,22)=6.3; p=0.007], showing significantly longer latency to lever press during the “reward/shock” trials [**: p-values<0.01] (Figure 1C). Similarly, we observed visual cue discrimination [F(2,22)=41.3; p<0.001], with decreased latency to enter the magazine when the cue signaled a reward [**: p-values<0.01] (Figure 1D). Additionally, rats exhibited differential latency to initiate the trials according to the previous outcome [F(2,22)=7.0; p=0.007], showing longer latency to initiate a new trial after experiencing a shock [**: p-values<0.01] (Figure 1B).

Together, these results indicate that the rats successfully learned the behavioral task and discriminated between reward, omission, and punishment trials.

### Single-unit electrophysiological recording

We recorded 106 BLA neurons with a population average firing rate of 19.3 ± 2.3 Hz. The average half-width of the action potentials was 0.68 ± 0.01 ms (Figure 1E, top panel). We recorded 111 BLA neurons with a population averaged firing rate of 4.4 ± 0.5 Hz. The average half-width of the action potentials was also 0.68 ± 0.01 ms. The average firing rate of both populations diverged from a normal distribution [Anderson Darling for CeA and BLA, respectively, A^2^=8 and 9.5; p-values<0.001] but not from log-normal distribution [A^2^=0.5 and 0.3; p=0.18 and p=0.52] (Figure 1E, bottom panel).

#### Amygdala neuronal activity during deliberation and execution processes

Z-scored firing rate of CeA and BLA significantly changed after the onset of auditory cues [time: CeA: F(63, 6615) = 15.6, p<0.001; BLA: F(62, 6820) = 10.5, p < 0.001, Figures 2A and 2B]. Only CeA neuronal activity discriminated between auditory cues [time x condition: CeA: F(126, 13230) = 2.3, p < 0.001; BLA: F(124, 13640) = 1.1, NS]. The late gradual increase in tonic CeA activity (1.4→5 s) significantly differed between cues [Friedman’s χ^2^(2)=18.4; p<0.0001], with reduced activity for “reward/shock” compared to “reward” and “reward/omission” cues [p <0.01] (Figure 2A insert). Conversely, neither the initial burst (0.5→1.3s) after post-cue onset nor the burst after lever presentation showed significant differences across conditions [Friedman’s χ^2^(2)=4.4 and χ^2^(2)<1; NS].

**Figure 2.**
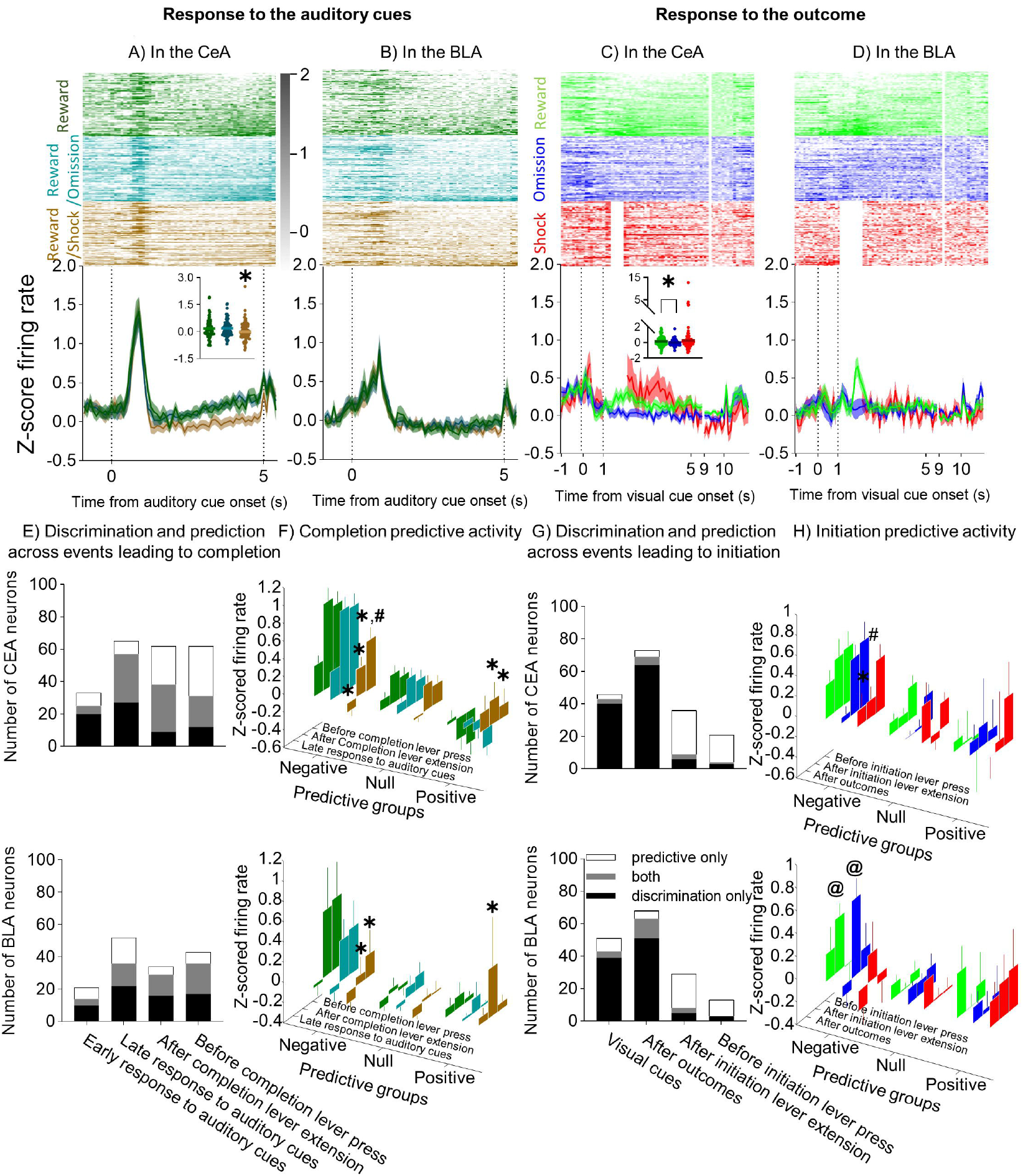
<u>(**A**) CeA and (**B**) BLA response to the auditory cues</u>: Heatmap of the individual neurons’ (top panels) and population mean (± SEM) of the z-scored firing rate (Hz) (bottom panels) of CeA and BLA neurons during 1 s preceding and the 5.4 s following the onset (0) of the auditory cue signaling “reward” (dark green line), “reward/omission” (teal line), or “reward/shock” (brown line) trials (n=106 and 111 neurons). In the heatmaps, individual neuron z-scored firing rates were stacked in ascending order according to their 2-second averaged activity across the 2 seconds following auditory reward cues, or their 3-s averaged activity following reward delivery. The insert illustrates the late CeA individual and average response (1.2 to 5 s after the onset) to the auditory cues *Significantly different from “reward/shock” trials, pairwise comparisons p<0.001. (<u>**C**) CeA and (**D**) BLA response to the outcomes</u>: Heatmap of the individual neurons’ (top panels) and population mean ± SEM of the z-scored firing rate (Hz) of CeA and BLA neurons across the 1 s preceding and the 5.5 s and across the 9 to 11 s following the onset (0) of the visual cues signaling reward (green line), omission of reward (blue line), or footshock punishment (red line) trials (n=106 and 111 neurons). The insert illustrates the CeA individual and average response to the outcome (4-s period following the offset of the visual cues), * Significantly different between omission of reward and reward, pairwise comparisons p<0.001. The top subpanels of panels (**E)** and (**G)** illustrates the number of CeA and BLA neurons which activity only discriminated between the different conditions (black), only predicting the latency to lever press on the completion lever (white) or simultaneously doing both (grey) across the different events leading to the completion of the trials. The bottom subpanels of panels (**E**) and (**G**) depict the same numbers across the different events leading to the initiation of new trials. (**F**) and (**H**) Changes in z-score firing rate of the CeA (top) or BLA (bottom) neural population negatively, positively, or not predicting the latency to completion lever pressing (**F**) or to initiation lever pressing (**H**) across conditions and across events. *: Significant differences between condition Benjamini-Hochberg’s corrected PLSD p<0.05, #: differences between events p<0.05 and @: difference between groups p<0.05.

Accordingly, the number of cue-discriminating CeA neurons peaked during the late tonic response phase (Figure 2E top panel). Approximately 60% of these neurons exhibited positive z-scores across events and conditions, except during the late response to the “reward/shock” auditory cue, where this proportion was inverted. Conversely, CeA neurons predicting response latency across conditions peaked after lever presentation, indicating partial overlap between cue-discriminating and response predicting neural populations. Out of the 92 CeA neurons that discriminated the cues or predicted latencies, 47 simultaneously did both.

In the BLA, the peak number of cue-discriminating neurons and latency-predicting BLA neurons occurred during the late auditory response (Figure 2E bottom panel). Among the 79 BLA neurons involved in cue discrimination or latency prediction, 36 performed both functions.

The power of neuronal activity to predict response latency determined across-condition extended to within the different conditions. Specifically, among the 84 CeA neurons and 68 BLA neurons that predicted latency variations across conditions at any point before completion lever press, a substantial proportion (58 and 42) simultaneously predicted latency fluctuations within specific conditions.

To examine how latency-predictive neuron activity relates to lever pressing, we categorized neurons by latency prediction (positive, negative, or none) and analyzed their z-scored activity across task events leading to lever pressing. In the CeA (Figure 2F, top panel), the activity of the predictive subpopulations significantly differed across conditions during the late auditory cue, lever presentation, and pre-lever press periods [group x conditions: F(4,206)=7.3, F(4,206)=15.9, F(4,206)=9.2, p-values<0.0001]. Neurons negatively predicting latency consistently exhibited greater activity in the “reward” compared to the “reward/shock” condition across all events [*: difference between conditions, p<0.001]. However, their response in the “reward/shock” condition progressively increased across events [condition x events: F(4,136)=4.3, p = 0.003], peaking just before lever pressing [# difference between events: p < 0.001]. In contrast, neurons positively predicting latency showed heightened activity in the “reward/shock” condition after lever presentation [condition x events: F(4,68)=2.6, p=0.04; * difference between conditions, p<0.001], maintaining a similar level of activity before lever pressing.

In the BLA (Figure 2F, bottom panel), predictive neuron activity diverged after lever presentation and before lever pressing [groups x events F(4,216)=5.7, F(4,216)=2.7, p<0.05]. Negatively predicting neurons showed sustained higher activity in “reward” vs. “reward/shock [*: difference between conditions: p-values <0.05], without changes across events [conditions x events F(4,84)=1.571, NS]. Positively predicting neurons exhibited greater activity in the “reward/shock” condition compared to the “reward” condition after lever presentation [*:difference between conditions, p=0.03] but not before lever pressing.

In summary, analysis of neuronal responses during deliberation and execution processes identified distinct amygdala neuron subpopulations, predictive of response latencies, exhibiting valence-specific responses, with a subset demonstrating dynamic activity, suggesting their roles in shaping decisions under varying reward and punishment.

#### Amygdala neuronal activity during reevaluation processes

CeA and BLA activity showed no difference across pre-cue and post-reward periods in rewarded trials from the different previous conditions (reward, reward/omission or reward/shock)[conditions, CeA: F(2, 210)<1 and BLA: F(2, 220)<1, NS]. Therefore, we pooled the z-score firing rates of all rewarded trials for further analysis.

Activity in both the CeA and BLA (Figures 2C and 2D), significantly changed after visual cues onset (- 1→1s) [time: F(19,1995)=2.9, F(19,2090)=2.2, p-values<0.01], during the late response (1.5→4.5s) to the different outcomes [F(29,3045)=2.1, F(29,3190)=1.7; p-values<0.01] and early response (0→1.5s) to reward and omission for BLA, but not CeA, (excluding shock at this stage because of recording artefact) [F(14,1540)=4.7, p<0.0001; F(14,1470)<1, NS]. CeA but not BLA discriminated between visual cues [time x conditions: F(38,3990)=1.9; p<0.001; F(38,4180)=1.1; NS], but no specific condition affected the averaged activity [Friedman’s χ^2^(2)=3.1; NS]. Outcome discrimination of CeA and BLA was more evident in early and late response to the outcome [CeA: F(14,1470)=2.3, F(58,6090)=1.8; BLA F(14,1540)=8.2; F(58,6380)=1.3; p-values<0.05]. with the averaged activity across the 4 seconds post-cue offset significantly greater in the CeA after reward than omission [Friedman’s χ^2^(2)=6.5; p=0.04; a: p=0.03] (Figure 2D insert) but no condition affected averaged activity in the BLA [χ^2^(2)=2; NS].

Unlike the pattern found during trial completion, CeA and BLA neurons showed temporal segregation of condition discrimination and trial initiation prediction (Figure 2G). Discrimination peaked after cue onset/offset, while the modest prediction peaked at lever presentation, with minimal overlap. Of the 91 neurons per region that discriminated conditions or predicted initiation latency, only a small subset performed both functions (12 CeA, 17 BLA).

BLA predictive neurons showed condition-specific activity after initiation lever presentation [interaction group x conditions: F(4,216)=2.9, p=0.02], driven by increased activity of neurons that negatively predicted latency to initiate a new trial following reward and omission [@: differences between groups, p-values<0.05]. However, these neurons did not show significant changes in activity across events [events x conditions: F(4,40) = 1.4, NS] (Figure 2H bottom panel). Conversely, CeA predictive neurons did not show significant differences across conditions during the events leading to initiation lever press but those that negatively predicted latency to initiate a new trial showed changes across events [F(4,92) = 2.6, p = 0.04]. After rats were shocked, activity before the lever press recovered from suppressed levels observed after the lever presentation [#: difference between events: p < 0.05; *: difference between conditions: p < 0.05] (Figure 2H top panel).

The similarity in neural dynamics between initiation and completion events suggests a shared neural population within the CeA. Indeed 17 of the 30 CeA neurons predicting initiation latency also predicted completion latency. However, 37 neurons were exclusively involved in completion, indicating increased CeA engagement during this phase.

CeA and BLA baseline firing rates weakly correlated with the determination coefficient for predicting trial completion latency from late auditory cue activity [ρ(104)=0.23; ρ(109)=0.22 p-values=0.02]. Additionally, baseline CeA activity weakly correlated with the effect size of the visual/auditory cues discrimination [ρ(104)=0.20; ρ(104)=0.28; p-values<0.05]. These results suggest a greater role of high-frequency neurons in stimulus discrimination and behavioral prediction.

## Discussion

In the present study, we extensively trained rats to ensure they effectively used the cues to guide their reward-seeking behavior. This resulted in faster latency to nosepoke when visual cues signaled an impending food reward. Conversely, rats exhibited longer latencies to lever-press to initiate a new trial following a probabilistic footshock, and took longer to complete a trial when auditory cues indicated the possibility of shock. The behavioral responses to probabilistic punishment were linked to differential neuronal activity patterns in the CeA and, to a lesser extent, the BLA.

Our findings confirm CeA neurons encode cues and outcomes valence. However, unlike Pavlovian fear conditioning studies reporting increased CeA activity to aversive cues (22), we observed population-wide reduction in CeA activity during “reward/shock” cues (15). For operant punishment conditioning as in the present study, aversive cues signal response-contingent outcomes, unlike Pavlovian cues that predict aversive events. This discrepancy suggests the CeA processes aversive cues differently across conditioning procedures.

We found no population-level discrimination of auditory cues in the BLA. Killcross et al. (10) reported that BLA lesions reversed punishment-induced suppression, while CeA lesions affect conditioned suppression manipulation (23). It is unlikely that suppression of operant responding in the present study relied on conditioned suppression, as the footshock presentation was contingent on lever pressing and was not directly paired with the auditory cue. The lack of population-level BLA punishment correlates also contrasts with Pavlovian conditioning studies showing BLA valence encoding (24-26), and with studies showing BLA discriminate between reward or punishment cues after 6 days of training in rats (27).

A possible explanation for the discrepancies between our findings and those of some prior studies is the extensive training our rats underwent. One challenge in studying punishment-induced suppression is to dissociate selective suppression of the operant punished responding from a generalized suppression of behavior (15). Another challenge in electrophysiological recording involves ensuring that rats continue to lever press despite potential punishment. To circumvent these challenges, we titrated the shock intensity for each rat so they would delay rather than stop responding during the “reward/shock” trials while maintaining consistent lever pressing in all other trial types. This required longer training (∼45 sessions) than previously reported in punishment studies. Such extended-training has been found to cause freezing behavior in the fear conditioning procedures to become insensitive to manipulations of BLA, while remaining sensitive to CeA manipulations (28,29). In addition, while the recruitment of dorsolateral striatum dopamine-dependent control over cocaine seeking is triggered by the BLA, its long-term maintenance depends instead on the CeA (30).

We speculate that overtraining in our study induced a shift in behavioral control from the BLA to the CeA, which consequently led to the observed absence of condition-specific neural responses in the BLA at a population level. This hypothesis fits the broader model that the BLA more generally mediates associative learning, supported by research on outcome devaluation and decision-making processes (31-34).

While population-level BLA activity showed no condition-specificity, individual neurons did. Their heterogeneous response (increased or decreased activity) may account for the discrepancy between our findings and those of Jean-Richard-dit-Bressel et al. (17) who reported increased population-level BLA activity during probabilistic punishment using fiber photometry, a method that obscures distinct neuronal subpopulations through signal averaging.

While most BLA and CeA neurons simultaneously distinguished the auditory cues and predicted trial completion, these functions were entirely segregated across events leading to the initiation of new trials. This indicates a greater role for the amygdala in translating contextual information into behavior during completion rather than initiation. This notion agrees with amygdala’s role in emotional processing and the observed greater arousal at initiation than at completion of a demanding task (35). We speculate that evaluating past experiences is less emotionally charged than anticipating future outcomes. Accordingly, we observed a greater proportion of predictive neurons, during task completion compared to initiation, especially in the CeA.

The mirroring of predictive neuronal activity dynamics across completion and initiation suggests the potential contribution of overlapping neurons involved in controlling responding under punishment threat. Overall, the failure to suppress instrumental responding in the face of punishment was associated with partial recovery from threat-induced inhibition of reward-seeking CeA neurons and the suppression of BLA punishment-sensitive neurons. These dynamic changes in neural activity appeared to counteract steady inhibition of punishment-associated CeA neurons and steady activation of reward-seeking BLA neurons.

The differential control of distinct neural populations over reward seeking under threat may explain the seemingly contradictory findings regarding the effect of CeA inactivation on punishment-induced behavioral suppression. While we and others have shown that inactivating the CeA reverses punishment-induced suppression of cocaine seeking (8,16), other studies have shown that CeA inactivation can enhance such suppression in punishment-resistant rats self-administering alcohol (18). It is therefore possible that depending on experimental variables (e.g., alcohol vs cocaine, punishment-resistant vs sensitive rats), responding under punishment might be under the control of the different amygdalar subpopulations of neurons.

In conclusion, we demonstrate that extensive training in punished reward-seeking responses is associated with specific neural activity patterns in the CeA and, to a lesser extent, the BLA. Notably, the failure to suppress reward seeking despite punishment appears to result from the dynamic interplay between the excitation of reward-driven and the inhibition of punishment-sensitive subpopulations of amygdala neurons. The present study lays the groundwork for characterizing the altered processes in CeA in pathological conditions such as drug addiction, where people with addiction continue to use drugs despite adverse consequences.

## Funding

The research was supported by the Intramural Research Program of NIDA (YS and GS) and a fellowship from the NIDA-INSERM program (YP).

## Authors contribution

YP designed and performed the experiments, analyzed the data, and wrote the paper with the other authors; BS helped with data analysis, the setup of the equipment, and the write up of the paper. AMT performed the experiments and helped with the writeup of the paper. DJD and EC helped with the write up of the paper, YS and GS were involved in different aspects of the experiments and the write-up of the paper.

## Competing Interests

The Authors have nothing to disclose.

## Notes

### Competing Interest Statement

The authors have declared no competing interest.

